# A host defense peptide mimetic, brilacidin, potentiates caspofungin antifungal activity against human pathogenic fungi

**DOI:** 10.1101/2022.09.28.509882

**Authors:** Thaila Fernanda dos Reis, Patrícia Alves de Castro, Rafael Wesley Bastos, Camila Figueiredo Pinzan, Pedro F. N. Souza, Suzanne Ackloo, Mohammad Anwar Hossain, David Harold Drewry, Sondus Alkhazraji, Ashraf S. Ibrahim, Hyunil Jo, William F. deGrado, Gustavo H. Goldman

## Abstract

*A. fumigatus* is the main etiological agent of a group of heterogeneous diseases called aspergillosis of which the most lethal form is the invasive pulmonary aspergillosis (IPA). Fungicidal azoles and amphotericin B are the first line defense against *A. fumigatus*, but fungistatic echinocandins, such as caspofungin (CAS), can be used as salvage therapy for IPA. Here, we screened repurposing libraries and identified several compounds that potentiate CAS activity against *A. fumigatus*, including the host defense peptide mimetic, brilacidin (BRI). BRI converts CAS into a fungicidal drug and potentiates voriconazole (VOR) against *A. fumigatus.* BRI increases the ability of both CAS and VOR to control *A. fumigatus* biofilm growth. BRI depolarizes the *A. fumigatus* cell membrane leading to disruption of membrane potential. By using a combination of protein kinase inhibitors and screening of a catalytic subunit null mutant library, we identified the mitogen activated protein kinase (MAPK) MpkA and the phosphatase calcineurin as mediators of the synergistic action of BRI. These results suggest the most likely BRI mechanism of action for CAS potentiation is the inhibition of *A. fumigatus* cell wall integrity (CWI) pathway. BRI potentiates CAS activity against *C. albicans*, *C. auris*, and *C. neoformans*. Interestingly, BRI overcomes the CAS-acquired resistance in both *A. fumigatus* and *C. albicans* and the CAS-intrinsic resistance in *C. neoformans*. BRI also has an additive effect on the activity of posaconazole (POSA) against several Mucorales fungi. Cell toxicity assays and fungal burden studies in an immunosuppressed murine model of IPA showed that BRI combined with CAS is not toxic to the cells and significantly clears *A. fumigatus* lung infection, respectively. Our results indicate that combinations of BRI and antifungal drugs in clinical use are likely to improve the treatment outcome of IPA and other fungal infections.

## Introduction

Fungal diseases occur in more than 1 billion people worldwide and are responsible for 1.5 million deaths (Bongomin et al., 2017; G. D. Brown et al., 2012). Aspergillosis encompasses a group of heterogeneous diseases caused by *Aspergillus spp.* (Rudramurthy *et al.,* 2019). In immunocompetent and immunosuppressed patients, aspergillosis is characterized by noninvasive and invasive diseases, respectively (Alastruey-Izquierdo et al., 2018; Denning et al., 2016; Patterson et al., 2016; Perlin et al., 2017; Rudramurthy et al., 2019). The most lethal form of aspergillosis in recipients of both hematopoietic stem cells and solid-organ transplants is invasive pulmonary aspergillosis (IPA) and *A. fumigatus* is the leading cause of this disease, which comprises more than 300,000 cases worldwide and is associated with a mortality rate of up to 90% in the most susceptible populations (Almyroudis et al., 2005; Azie et al., 2012; G. D. Brown et al., 2012; Gonçalves et al., 2016; Guinea et al., 2010; Rudramurthy et al., 2019; Rüping et al., 2008).

Azoles (itraconazole, posaconazole, voriconazole, and isavuconazole) are fungicidal drugs for *A. fumigatus* and are used as first-line therapy against IPA while the fungistatic echinocandins, such as caspofungin (CAS), can be used as salvage therapy and have been recommended in combination therapies against emerging azole-resistant infections (Jenks & Hoenigl, 2018; Mavridou et al., 2015; Ostrosky-Zeichner & Al-Obaidi, 2017). Azoles inhibit the ergosterol biosynthesis pathway by directly targeting the *cyp51/erg11* encoding the lanosterol 14-demethylase (Perfect, 2017; Robbins et al., 2017). CAS acts by noncompetitively inhibiting the fungal β-1,3-glucan synthase (Fks1), required for the biosynthesis of β-1,3-glucan, and essentially blocking fungal cell wall synthesis (Perlin, 2015). Considering the paucity of available antifungal drugs and the increasing number of azole-resistant environmental isolates, clinical azole-resistant *A. fumigatus* isolates are currently a crucial problem and a major threat to immunosuppressed patients (Arikan-Akdagli et al., 2018; Chen et al., 2020; Garcia-Rubio et al., 2017; Resendiz Sharpe et al., 2018; Verweij et al., 2016; Wiederhold, 2017; Wiederhold & Verweij, 2020; Bastos *et al*., 2021).

Because of the scarcity in antifungal agents currently in development (Hoenigl *et al*., 2021), repurposing of currently approved drugs alone or in combination with currently used antifungal agents, presents a potential opportunity for the discovery of new antifungal agents (Nosengo, 2016; Kaul *et al*., 2019; Iyer *et al*., 2021). By using this strategy, several compounds have already been identified as potential new antifungal agents, and more importantly as potentiators of antifungal drugs currently in clinical use (Rhein *et al*., 2016; Joffe *et al*., 2017; Duffy *et al*., 2017; Wall *et al*., 2019; Revie *et al*., 2020; Iyer *et al*., 2020, 2021). Here, we screened four chemical collections containing a total of 1,402 compounds aiming to identify compounds that could enhance the *in vitro* CAS activity against *A. fumigatus.* We identified 6 CAS enhancers (compounds that can partially inhibit the fungal growth at 20 µM and have increased inhibition when combined with CAS) and 12 CAS synergizers (compounds that cannot inhibit the fungal growth at 20 µM but have increased inhibition when combined with CAS). Among the CAS synergizers, we closely investigated a small molecule host defense peptide mimetic, brilacidin (BRI), which is in clinical development for multiple indications. BRI has previously exhibited broad-spectrum inhibitory activity in bacteria and viruses, as well as immunomodulatory/anti-inflammatory properties (Innovation Pharmaceuticals Inc., 2022, http://www.ipharminc.com/brilacidin-1). BRI is able to synergize with echinocandin and azoles activities converting CAS into a fungicidal drug and enhancing the cidal activity of voriconazole (VOR) and posaconazole (POSA) against *A. fumigatus* and Mucorales fungi, respectively. Further, BRI is efficient against *A. fumigatus* biolfilm and can overcome CAS-resistance but not VOR-resistance. BRI can also synergize with CAS against other human pathogenic fungi, such as *C. albicans, C. auris*, and *C. neoformans*. By a combination of screening an *A. fumigatus* phosphatase null mutant library and a collection of protein kinase inhibitors, we were able to assign the CAS synergism mechanism of action to the cell wall integrity (CWI) pathway. Both calcineurin and the CWI mitogen-activated protein kinase (MAPK) MpkA are important for the mechanism of action of BRI in increasing CAS activity.

## Results

### Screening of the COVID Box, Pandemic Response Box, NIH Clinical, and epigenetic compound libraries

To identify compounds that can enhance or synergize with CAS activity against *A. fumigatus*, we used the Minimal Effective Concentration (MEC) assay to screen the fungus susceptibility to four chemical drug libraries: (i) the COVID Box (containing 160 compounds, see https://www.mmv.org/mmv-open/archived-projects/covid-box); (ii) the Pandemic Response Box (containing 400 compounds, see https://www.mmv.org/mmv-open/pandemic-response-box/about-pandemic-response-box); (iii) the National Institutes of Health (NIH) clinical collection (NCC) (containing 727 compounds; see https://pubchem.ncbi.nlm.nih.gov/source/NIH%20Clinical%20Collection); and (iv) the epigenetic probe library (containing 115 compounds, see https://www.sgc-ffm.uni-frankfurt.de/). In total, two methods were employed to assess 1,402 compounds by using a combination of 0.2 µg/ml of CAS (the minimum effective concentration, MEC) and up to 20 µM of each compound compared to the effect on growth of *A. fumigatus* of each drug alone. First, we assessed growth by two independent rounds of visual inspection and selected 17 compounds that could inhibit *A. fumigatus* growth. Second, *A. fumigatus* growth in the presence of CAS 0.2 µg/ml alone, each of these 17 compounds at 20 µM alone, and a combination of each of these compounds from 0.6 to 20 µM plus CAS 0.2 µg/ml was quantified by using Alamar blue (Figure 1A). Based on this assay we defined enhancers as the compounds that alone could inhibit over 30 % of *A. fumigatus* metabolic activity but in combination with CAS inhibited even more, while synergizers were defined as compounds which alone inhibited less than 30 % of the fungal metabolic activity but in combination with CAS inhibited more than 30 %. Five compounds were classified as enhancers (Figure 1A): (i) chlormidazole (https://go.drugbank.com/drugs/DB13611) and (ii) ravuconazole (https://go.drugbank.com/drugs/DB06440; both inhibiting ergosterol biosynthesis), (iii) 5-fluorocytosin (https://go.drugbank.com/unearth/q?c=_score&d=down&query=5-fluorocytosin&searcher=drugs; that inhibits the RNA and DNA biosynthesis), (iv) ciclopyxox (https://go.drugbank.com/unearth/q?utf8=%E2%9C%93&searcher=drugs&query=ciclopyxox; it is thought to act through the chelation of polyvalent metal cations, such as Fe^3+^ and Al^3+^; Niewerth *et al*., 2003), and (v) MMV1593544 (a possible antiviral compound that inhibits SARS-CoV-2 infection *in vitro*; Holwerda *et al*., 2020). Twelve compounds were classified as synergizers (Figures 1A and 1B): (i) toremifene (https://go.drugbank.com/drugs/DB00539; a nonsteroidal triphenylethylene derivative used as an antitumor drug that appears to bind to the estrogen receptors competing with estradiol), (ii) brilacidin (https://go.drugbank.com/drugs/DB12997; a compound that acts as a mimetic of host defense peptides; Mensa *et al*., 2014), (iii) MMV1634399 (a quinoline anti-malarial; Reader *et al*, 2021), (iv) Diiodoemodin or MMV1581545 (an anti-bacterial emodin derivative; Ji *et al*., 2020), (v) PPTN (a potent, high-affinity, competitive and highly selective nucleotide-sugar-activated P2Y14 receptor antagonist; Barrett *et al*., 2013), (vi) triclopidine (a prodrug that is metabolised to an active form, which blocks the ADP receptor that is involved in GPIIb/IIIa receptor activation leading to platelet aggregation; https://go.drugbank.com/drugs/DB00208), (vii) loxoprofen (a non-steroidal anti-inflammatory drug that acts as a non-selective inhibitor of cyclooxygenase enzymes, which are responsible for the formation of various biologically active pain, fever, and inflammatory mediators; https://go.drugbank.com/drugs/DB09212), (viii) regorafenib (a small molecule inhibitor of multiple membrane-bound and intracellular kinases involved in normal cellular functions and in pathologic processes such as oncogenesis, tumor angiogenesis, and maintenance of the tumor microenvironment; https://go.drugbank.com/drugs/DB08896), (ix) OSU 03012 (a potent inhibitor of recombinant phosphoinositide-dependent kinase 1; Tseng *et al*., 2005), (x) MMV1782211 (an inhibitor of the SARS-CoV-2 main protease; https://chemrxiv.org/engage/chemrxiv/article-details/60c753ed469df403bef44e65), (xi) MMV1782350 and (xii) MMV1782097 (two uncharacterized antivirals).

**Figure 1.**
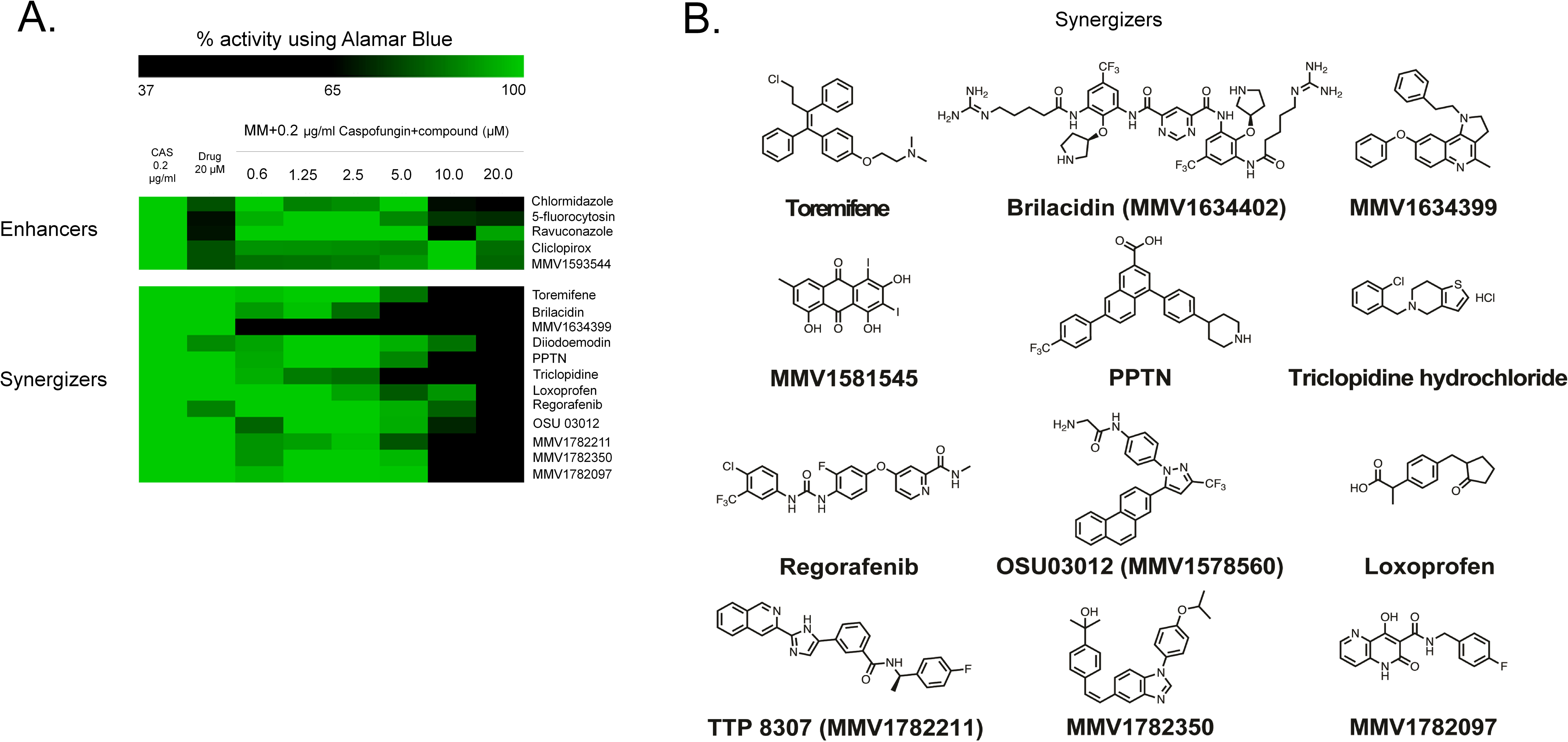
Screening of repurposing chemical libraries identify several compounds that enhance or synergize caspofungin activity. (A) Heat map of % of activity using Alamar blue. The % of activity is based on *A. fumigatus* grown in the absence or presence of a specific compound (MM+CAS 0.2 µg/ml or enhancers or synergizers 20 µM alone, or a combination of enhancer or synergizers from 0.6 to 20 µM) divided by the control (MM), both grown for 48h at 37°C. Hierarchical clustering was performed in Multiple Experiment Viewer (MeV) (http://mev.tm4.org/), using Pearson correlation with complete linkage clustering. Heat map scale and gene identities are shown. (B) Chemical structures of the synergizers.

Taken together, these results suggest we were able to identify many compounds that can enhance or synergize the activity of CAS against *A. fumigatus*. The synergizers have very different mechanisms of action and targets, most of them apparently not conserved in *A. fumigatus*.

### BRI converts CAS into a fungicidal drug and overcomes CAS-resistance

We decided to concentrate our further analysis on the host defense peptide mimetic brilacidin (BRI) since this drug candidate has undergone several clinical trials showing therapeutic benefit. BRI MIC for *A. fumigatus* is 80 µM (Table 1) and the *A. fumigatus* conidial viability was tested after 48 h of exposure to a combination of CAS 0.2 or 0.5 µg/ml combined with 20 µM BRI (Figure 2A). The combination of 0.2 or 0.5 µg/ml of CAS with 20 µM BRI reduced *A. fumigatus* conidial viability by 85% and 100%, respectively (Figure 2A). BRI 20 µM could also synergize with subinhibitory VOR concentrations of 0.125 and 0.25 µg/ml by reducing the *A. fumigatus* conidial viability by 92% and 99%, respectively (Figure 2A).

**Figure 2.**
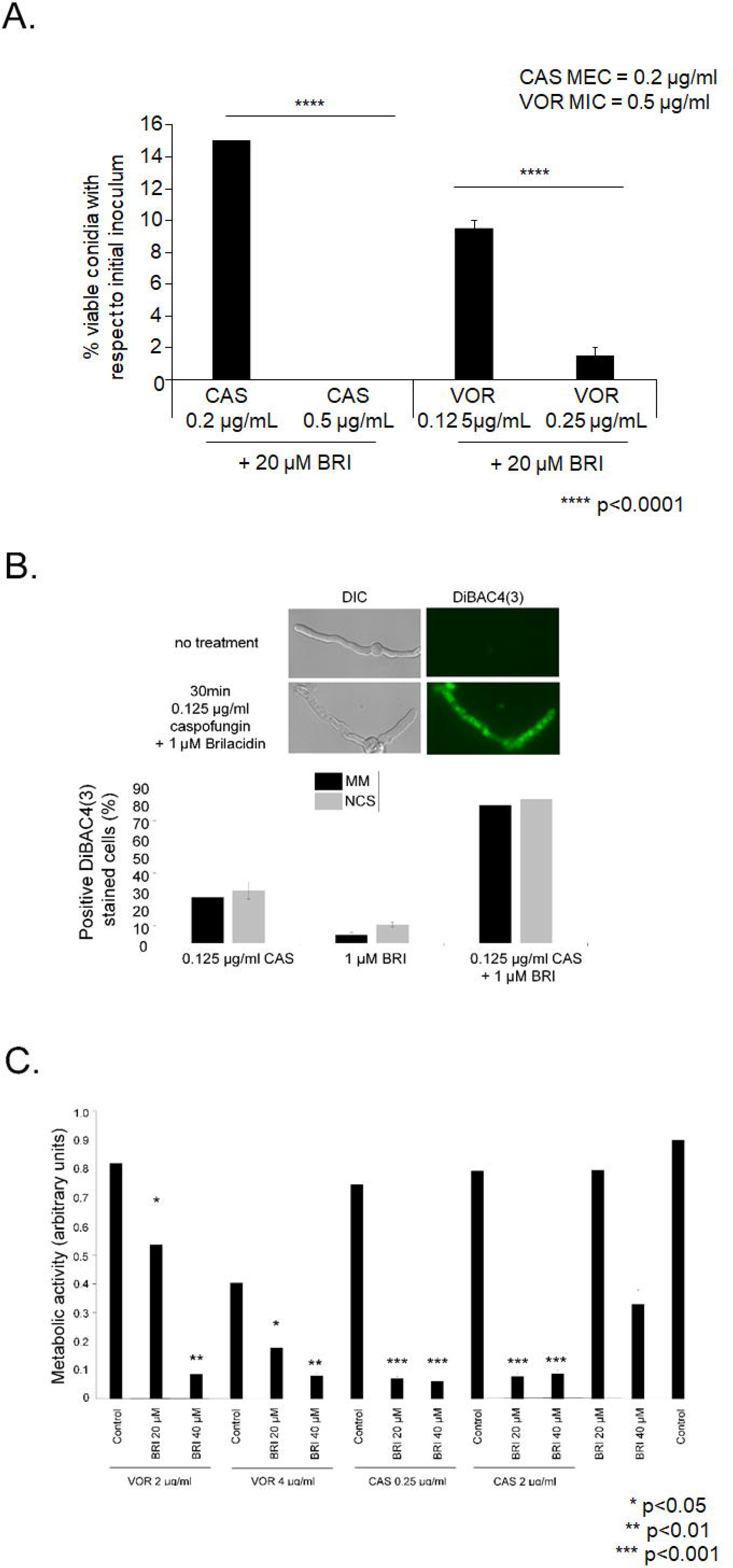
BRI can convert CAS into a fungicidal drug. (A) *A. fumigatus* conidia were incubated for 48 hs at 37°C with different combinations of BRI+CAS and BRI+VOR. After this period, non-germinated conidia were plated on MM and colony forming units (CFUs) were assessed. The results are expressed as the % of viable conidia with respect to initial inoculum and are the average of three repetitions ± standard deviation (*p* < 0.0001). (B) BRI+CAS disrupts the *A. fumigatus* membrane potential. *A. fumigatus* was grown for 16h at 37°C and exposed to CAS 0.125 µg/ml, BRI 1 µM or CAS 0.125 µg/ml+BRI 1 µM for 30 min and 3 µg/ml DIBAC_4_(3). Germlings were previously transferred to MM without glucose (non-carbon source, NCS) for 4h or not (MM) before adding CAS, BRI, or CAS+BRI for 30 min and subsequently DIBAC_4_(3). The results are expressed as the % of fluorescent cells and are the average of three repetitions of 50 germlings each ± standard deviation. (C) Metabolic activity expressed by XTT of *A. fumigatus* biolfilm formation in the presence of VOR or CAS alone and combinations of BRI+VOR and BRI+CAS. Biofilm was formed for 24h of incubation at 37°C and after this period 50µL of fresh MM containing CAS, VOR or a combination of VOR+CAS or CAS+BRI were added to the biofilm to reach the final concentration as indicated and incubated for further 12h at 37°C. Untreated biofilm was used as a positive control. The XTT assays were performed in six replicates and the results are expressed as the average ± standard deviation (*, *p* < 0.05, **, *p <* 0.01, ***, *p* < 0.001, and ****, *p* < 0.0001).

**Table 1.**
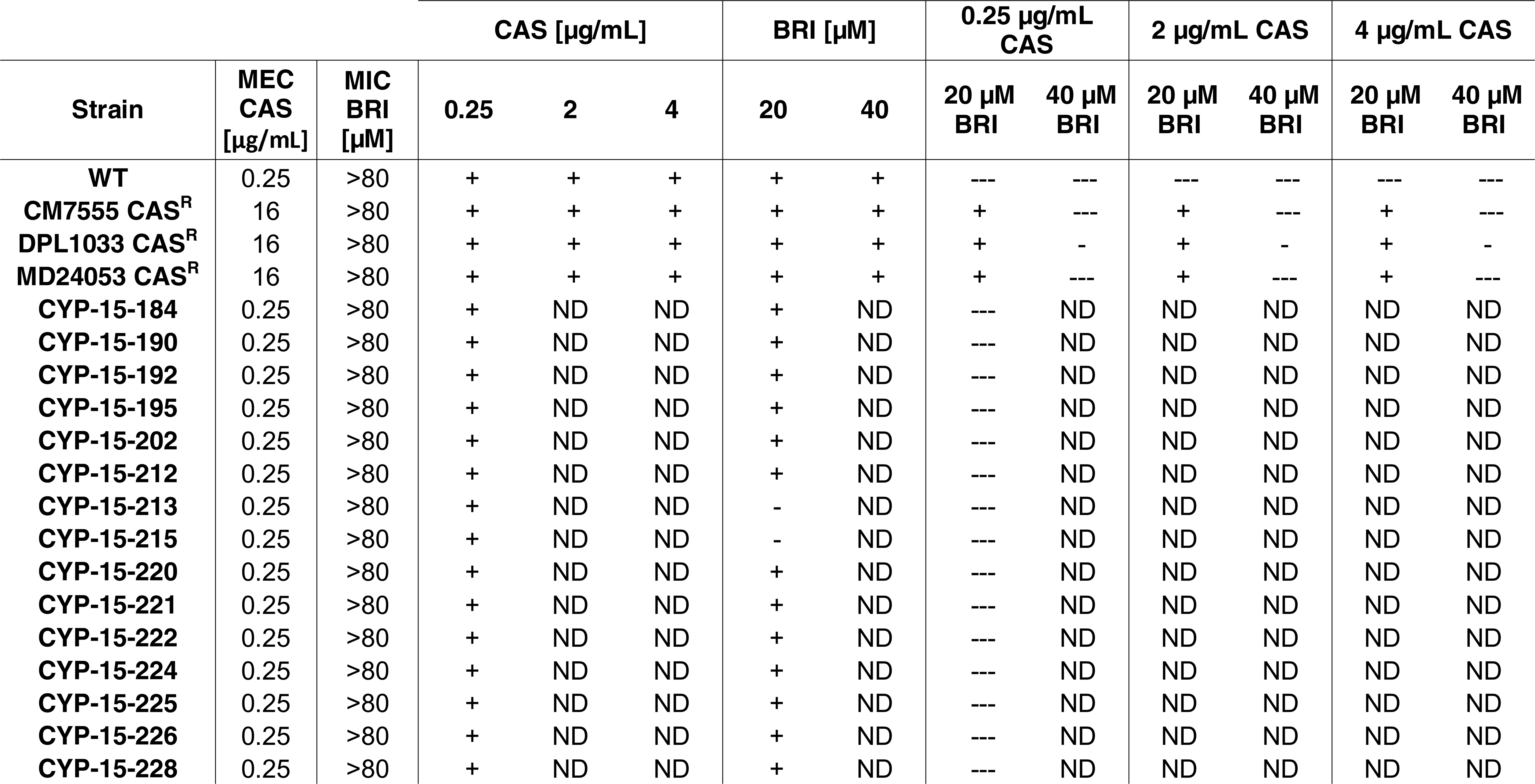
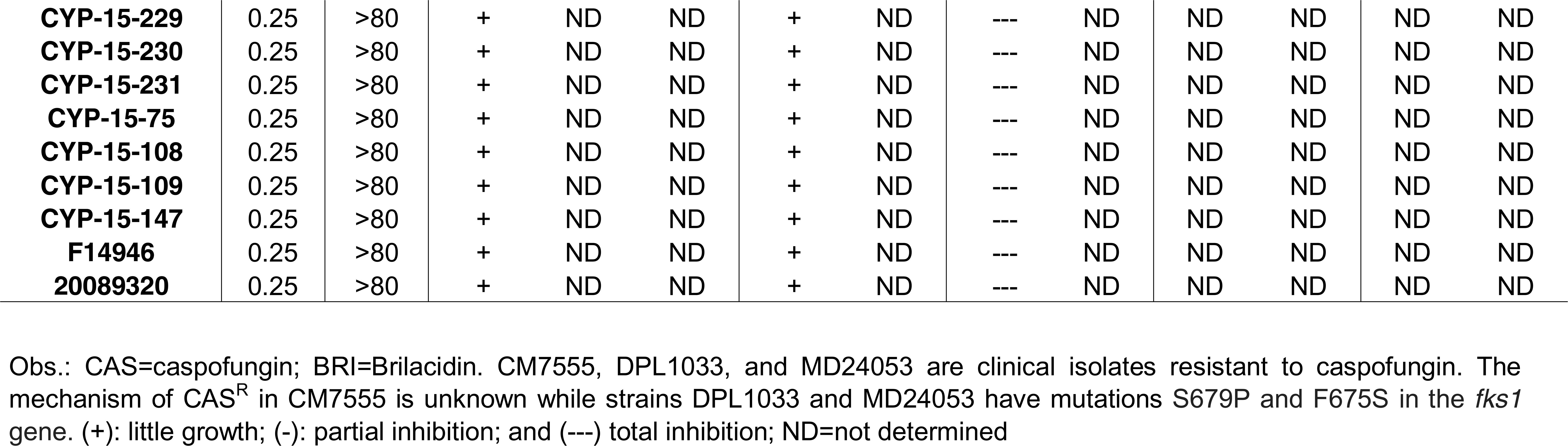
BRI+CAS can overcome *A. fumigatus* CAS-resistance

Antimicrobial peptides target directly or indirectly the microorganism plasma membrane disrupting their membrane potential (Lima *et al*., 2021; Veerana *et al*., 2021), and BRI acts by a similar mechanism (Mensa *et al*., 2014; Tew *et al*., 2002, 2010). We determined the effect of BRI+CAS on the resting membrane potential by using the fluorescent voltage reporter DIBAC_4_(3) (increase in the fluorescent intensity indicates membrane depolarization; Figure 2B). Untreated *A. fumigatus* had 0% fluorescent germlings while exposure to either 0.125 µg/ml CAS, 1 µM BRI, or a combination of both resulted in ∼25%, 5%, and 80% fluorescent germlings, respectively (Figure 2B). Germlings previously transferred to (minimal medium) MM without glucose (non-carbon source, NCS) for 4 hours before adding CAS, BRI, or CAS+BRI for 30 min and subsequently DIBAC_4_(3) have not shown any fluorescent difference with the cells that were grown in the presence of the carbon source glucose (Figure 2B). These results suggest that BRI as well as CAS is passively transported through the cell membrane without the need for ATP.

*A. fumigatus* biofilm formation is more resistant to antifungal agents (Morelli *et al*., 2021). BRI alone at 20 or 40 µM can inhibit about 10 and 60% biofilm formation, respectively (Figure 2C) while VOR alone at 2 µg/ml or 4 µg/ml were able to inhibit about 10 and 50% biofilm formation (Figure 2C); CAS alone at 0.25 µg/ml or 2.0 µg/ml were able to inhibit about 10% biofilm formation (Figure 2C). The different combinations BRI+VOR or BRI+CAS were able to potentiate the inhibition of the biofilm formation about 50 to 90% (Figure 2C).

We also evaluated if the combination of BRI+CAS can inhibit CAS-resistant and VOR-resistant *A. fumigatus* clinical isolates (Table 1). Specifically, we tested a range of 0.25 to 4 µg/ml of CAS with BRI at 20 and 40 µM and VOR at concentrations of 0.5 and 2 µg/ml with BRI 20 and 40 µM. BRI had no activity against 25 *A. fumigatus* clinical isolates susceptible to CAS (MEC CAS of 0.25 µg/ml) and 3 CAS-resistant clinical strains [MEC CAS of 16 µg/ml; strains DPL1033, and MD24053 with known *fks1* mutations; and strain CM7555 with an unknown mutation(s)]. Interestingly, addition of BRI at 20 or 40 µM to CAS either partially or completely inhibited the growth of all tested strains including those that are resistant to CAS or with known resistance to azoles (Table 1). Thus, BRI clearly potentiates CAS activity against CAS- or VOR-resistant strains of *A. fumigatus*.

In contrast to the potentiation of CAS activity by BRI, addition of BRI (at 20 or 40 µM) to VOR had no effect on the resistant nature of 22 clinical isolates [15 strains with the TR34/L98H mutation and 7 strains with unknown mutation(s)] to VOR (Supplementary Table S1). In the experiment, effects were observed for 2 clinical isolates: one strain (CYP15-15-109) was totally inhibited by VOR, showing wild-type response; and, one other strain (CYP15-15-147) was partially inhibited by VOR 0.5 µg/ml+BRI but the inhibition was not seen at VOR 2.0 µg/ml+BRI (Supplementary Table S1). Curiously, the VOR-resistant clinical isolates were not inhibited by a combination of BRI+VOR but they were inhibited by BRI+CAS (compare Table 1 with Supplementary Table S1). Most of the VOR-resistant strains have increased accumulation of ergosterol since the tandem-repeat mutations at the promoter region increase the *erg11A* expression and consequently the ergosterol production (Hagiwara *et al*., 2016). Ergosterol is essential for the integrity and fluidity of fungal cell membranes and azole-induced depletion of ergosterol alters the membrane sterol composition, its stability and arrests fungal growth (Shapiro *et al*., 2011).

Taken together, these results indicate that the combination CAS+BRI can depolarize the cell membrane converting CAS from a fungistatic into a fungicidal drug for *A. fumigatus*. BRI+CAS can decrease *A. fumigatus* biofilm formation and completely or partially overcome CAS-resistance in echinocandin-resistant *A. fumigatus* clinical isolates. VOR-resistant isolates are sensitive to CAS+BRI combinations but most of them are not sensitive to BRI+VOR combinations.

### BRI is impacting *A. fumigatus* calcineurin signaling and the cell wall integrity (CWI) pathway

To assess the mechanism of action of BRI, a collection of 58 protein kinase inhibitors (PKI, at a concentration of 20 µM; Supplementary Table S2) was screened for *A. fumigatus* growth and corresponding metabolic activity alone or together with 20 µM BRI (Supplementary Table S2). Two PKIs, a p21-Activated Kinase Inhibitor FRAX486 and a STK25 inhibitor PP121, both members of the sterile 20 superfamily of kinases, are identified as potentiating the BRI activity against *A. fumigatus* (Figures 3A and 3B). The p21 activated kinases (PAKs) belong to the family of Ste20-related kinases and these kinases have been shown to be involved in signaling through mitogen activated protein kinase (MAPK) pathways (Boyce and Andrianopoulos, 2011). The closest STK25 homologues are cAMP-mediated signaling proteins Sok1p in *Saccharomyces cerevisiae* whose overexpression suppresses the growth defect of mutants lacking protein kinase A activity (Ward *et al*., 1994). We also screened a library of 25 *A. fumigatus* null mutants for phosphatase catalytic subunits (Winkelströter *et al*., 2015) for sensitivity to BRI 20 µM. We identified a single phosphatase mutant, Δ*calA* (*calA* encodes the calcineurin catalytic subunit), as more sensitive to BRI 20 µM (Supplementary Table S3).

**Figure 3.**
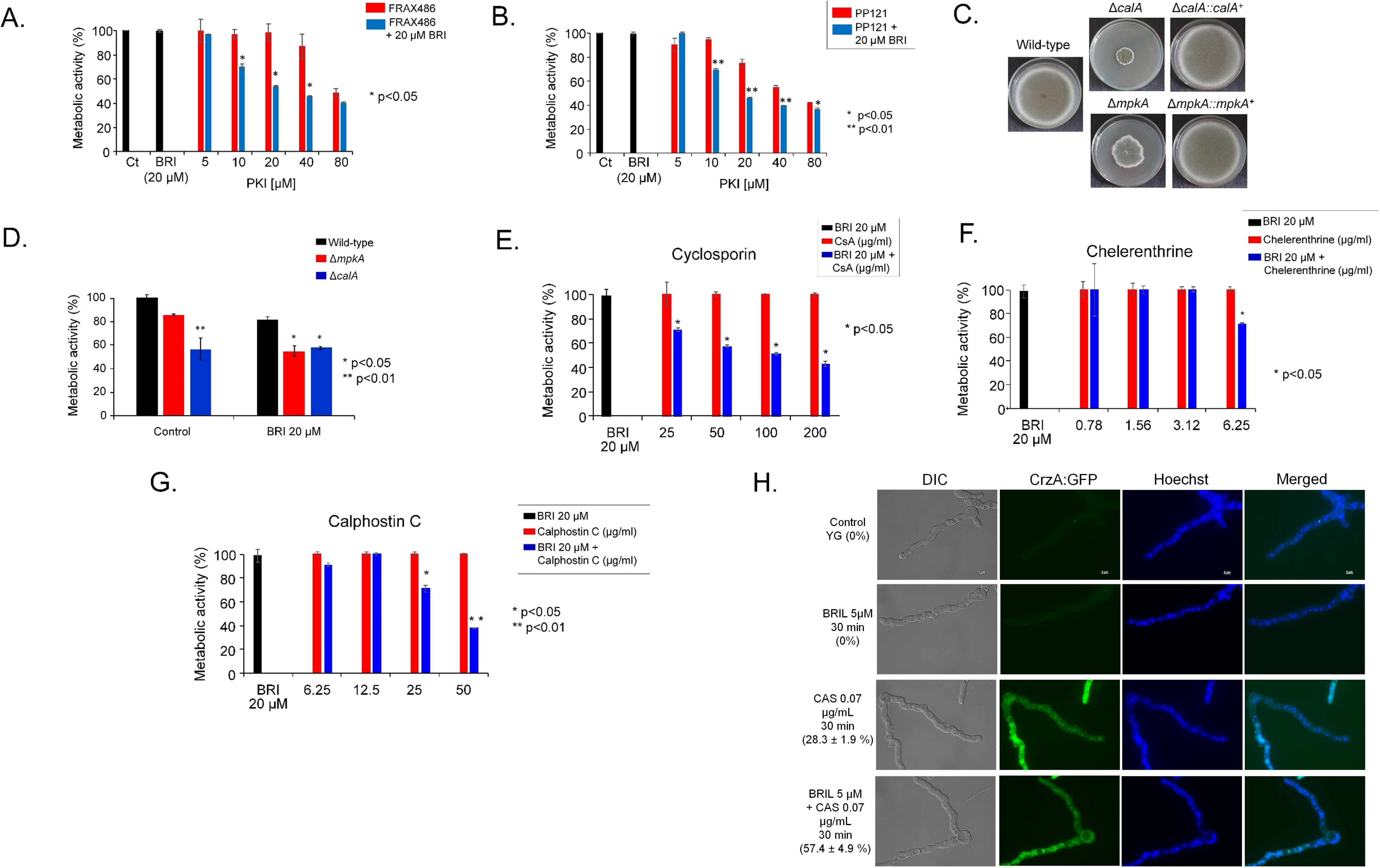
Calcineurin and the MAPK MpkA are important for the BRI+CAS synergism. (A) Metabolic activity expressed by Alamar blue of *A. fumigatus* grown for 48 hours in the absence or presence of BRI 20 µM or BRI 20 µM+FRAX486 5 to 80 µM (*, *p* < 0.05). (B) Metabolic activity expressed by Alamar blue of *A. fumigatus* grown for 48 hours in the absence or presence of BRI 20 µM or BRI 20 µM+PP121 5 to 80 µM (*, *p* < 0.05 and **, *p* < 0.01). (C) Growth of the wild-type, Δ*calA*, Δ*calA::calA^+^*, Δ*mpkA*, and Δ*mpkA::mpkA^+^* on MM for 5 days at 37°C. (D) Metabolic activity expressed by Alamar blue of *A. fumigatus* wild-type, Δ*calA*, and Δ*mpkA* grown for 48 hours in the absence or presence of BRI 20 µM (*, *p* < 0.05 and **, *p* < 0.01). (E) Metabolic activity expressed by Alamar blue of *A. fumigatus* grown for 48 hours in the absence or presence of BRI 20 µM or BRI 20 µM+cyclosporin 25 to 200 µg/ml (*, *p* < 0.05). (F) Metabolic activity expressed by Alamar blue of *A. fumigatus* grown for 48 hours in the absence or presence of BRI 20 µM or BRI 20 µM+Chelerenthrine 0.78 to 6.25 µg/ml (*, *p* < 0.05). (G) Metabolic activity expressed by Alamar blue of *A. fumigatus* grown for 48 hours in the absence or presence of BRI 20 µM or BRI 20 µM+Calphostin C 6.25 to 50 µg/ml (*, *p* < 0.05 and **, *p* < 0.01). All the results with Alamar blue are the average of three repetitions ± standard deviation. (H) CrzA:GFP translocation to the nucleus when germlings were exposed to BRI 5 µM, CAS 0.07 µg/ml or BRI 5 µM +CAS 0.07 µg/ml. Conidia were germinated in YG medium and grown for 13 hours and then exposed or not to BRI, CAS, and BRI+CAS for 30min. The results are the average of two repetitions with 30 germlings each ± standard deviation.

Considering the importance of calcineurin in the signaling response to *A. fumigatus* osmotic stress and the cell wall integrity pathway (da Silva Ferreira *et al*., 2007; Ries *et al*., 2017; de Castro *et al*., 2014, 2019) and the fact that PAKs have been shown to be involved in signaling through MAPK pathways (Boyce and Andrianopoulos, 2011), we tested the BRI growth inhibition of four *A. fumigatus* null mutants for MAPK Δ*sakA*, Δ*mpkC*, Δ*mpkB* and Δ*mpkA* which regulate osmotic stress and CWI pathways responses (Brown and Goldman, 2016; Yaakoub *et al*., 2021). We identified only Δ*mpkA* as being more sensitive to BRI 20 µM (Supplementary Table S3). Both mutants Δ*calA* and Δ*mpkA* have severe growth defects (Figure 3C) and further validation by using Alamar blue metabolic activity assay showed that Δ*calA* strain had similar metabolic activity in the presence or absence of BRI (Figure 3D). In contrast, the Δ*mpkA* mutant had a significantly decreased metabolic activity when grown in the presence of BRI (Figure 3D). Hsp90 chaperone is important for the regulation of several fungal signaling pathways either by physical interaction or regulation of their expression (Leach *et al*., 2012). *A. fumigatus* calcineurin and CWI kinases are possible Hsp90 client proteins (Lamoth *et al*., 2012, 2013, 2016). We investigate the influence of Hsp90 on BRI activity by determining the relationship between the Hsp90 inhibitor geldanamycin (GEL) and BRI in a checkerboard assay for which GEL concentrations of 0 to 25 µg/ml and BRI concentrations of 0 to 80 µM were used. The Fractional Inhibitory Concentration (FIC) index for these two drugs was 0.64 indicating additive effect for GEL when added to BRI against *A. fumigatus* (Supplementary Figure S1).

Cyclosporine (CsA) is a specific inhibitor of calcineurin and there is synergy, in inhibition of *A. fumigatus* metabolic activity, between increasing concentrations of CsA and BRI 20 µM (Figure 3E). *A. fumigatus* protein kinase C (PKC) is important for the activation of the CWI pathway (Rocha *et al*., 2015) and PKC inhibitors such as chelerenthrine and calphostin C also synergize with BRI 20 µM (Figures 3F and 3G). *A. fumigatus* calcineurin regulates the activity of the CrzA transcription factor with the phosphorylated form accumulating in the cell cytosol, and in response to several stimuli, including CAS exposure, calcineurin dephosphorylates CrzA leading to its re-localization to the nucleus (Ries *et al*., 2017; de Castro *et al*., 2014, 2019). When a functional CrzA:GFP strain is not exposed to any drug, or exposed to BRI (5 µM), or to a sub-inhibitory concentration of CAS (0.07 µg/ml), 0, 0 and 28.3 %, respectively, of the germlings have CrzA:GFP in the nuclei, while 57.4 % are in the nuclei when this strain is exposed to a combination of BRI+CAS (Figure 3H). There results strongly suggest that CrzA is important for the synergistic activity of BRI+CAS.

Taken together, these results strongly indicate that CalA/CrzA and MpkA are important for BRI activity, and BRI is most likely impacting the *A. fumigatus* CWI pathway.

### BRI can potentiate caspofungin activity in *C. neoformans*, *C. albicans*, and *C. auris*

We also investigated if BRI could potentiate CAS activity in other human fungal pathogens, such as *C. neoformans*, *C. albicans*, and *C. auris*. The MICs for BRI in *C. neoformans*, *C. albicans*, and *C. auris* are 2.5 µM, 80 µM and 80 µM, respectively (Table 2). CAS lacks significant activity against *C. neoformans* (Johnson and Perfect, 2003) and only high CAS concentrations, such as CAS 32 µg/ml can completely inhibit *C. neoformans* metabolic activity (as determined by XTT) and CAS 16 µg/ml can decrease survival (colony forming units, CFUs) by about 50 % (Figure 4A). BRI 0.625 µM (0.25xMIC) potentiates CAS activity (0.25 to 0.5 µg/ml of CAS) resulting in complete inhibition of *C. neoformans* metabolic activity and growth (Figure 4A). Similarly, complete inhibition of *C. albicans* metabolic activity (XTT) and survival (CFUs) shifted from a concentration of CAS at ∼ 0.125 µg/ml in the absence of BRI to a concentration of 0.015 µg/ml of CAS when 20 µM of BRI is added (*i.e.* an 8-fold reduction in the MIC of CAS) (Figure 4B). BRI also partially suppressed the CAS-resistance of *C. albicans* CAS-resistant clinical isolates (Figure 4C). BRI MICs for each *C. albicans* CAS-resistant isolate is 20 to 80 µM (Table 2) and the combination of BRI 5 to 20 µM (0.25x MIC)+CAS 0.5 µg/ml decreased CAS-resistance of DPL1006, DPL1007, DPL1009, DPL1010, and DPL1011 7-, 10-, 4-, 10-, and 2-fold, respectively (Figure 4C). Finally, BRI MIC for *C. auris* is 80 µM (Table 2) and BRI 10 µM + CAS 0.125 µg/ml inhibited about 95 % *C. auris* metabolic activity in clinical isolates 467/2015, 468/2015, 469/2015, 470/2015, and 474/2015 (Figure 4D). In one of these clinical isolates, 467/2015, BRI 10 µM+CAS 0.5 µg/ml is able to inhibit 100 % metabolic activity and survival, potentiating caspofungin activity by at least 2-fold (Figure 4D).

**Figure 4.**
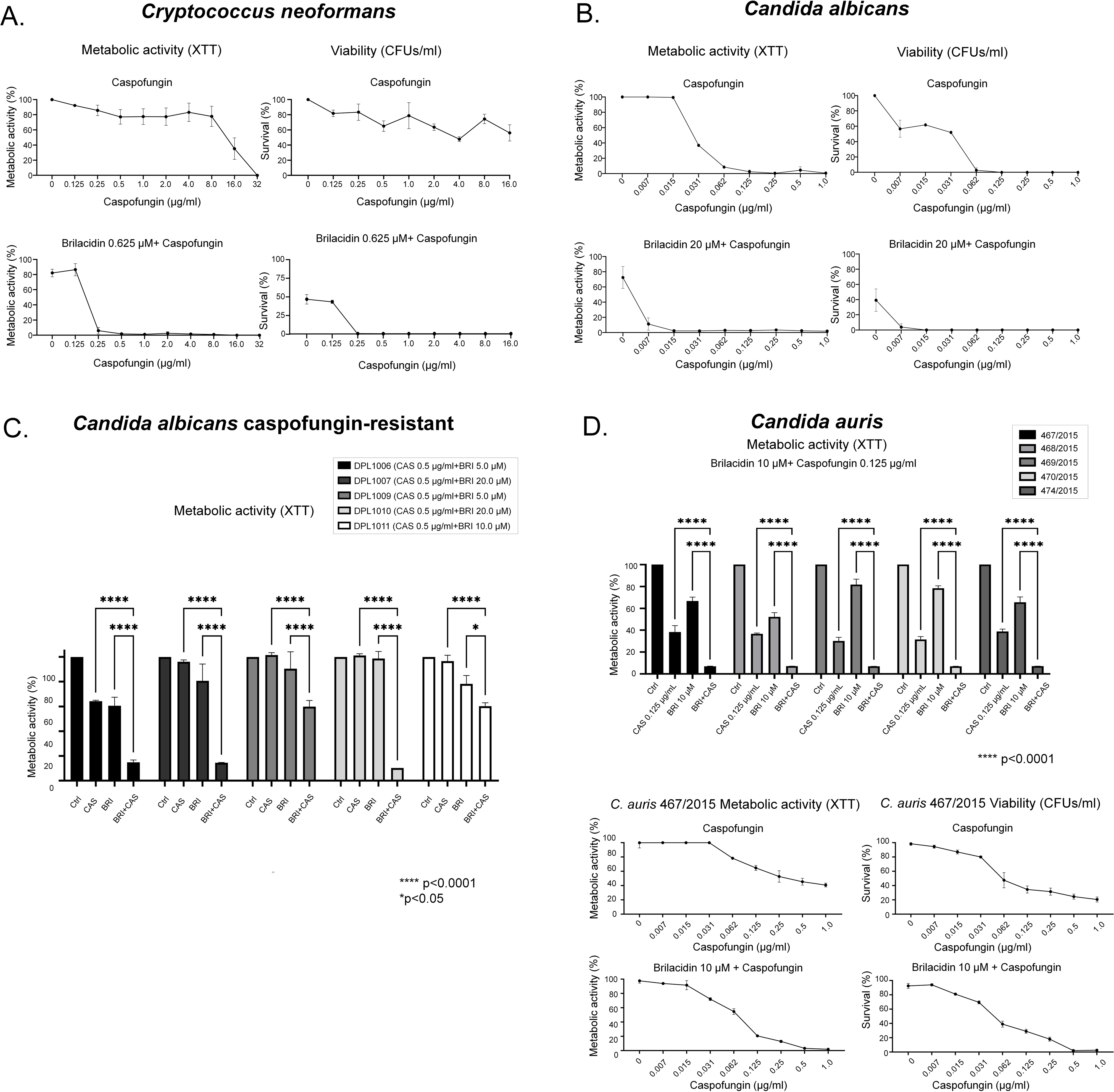
BRI can synergize CAS in other human fungal pathogens. (A) Metabolic activity expressed by XTT of *C. neoformans* grown for 48 hours in the absence or presence of CAS 0 to 32 µg/ml or BRI 0.625 µM+CAS 0 to 32 µg/ml. Percentage of survival expressed as colony forming units/ml of *C. neoformans* cells grown for 48 hs in the absence or presence of CAS 0 to 16 µg/ml or BRI 0.625 µM+CAS 0 to 16 µg/ml. (B) Metabolic activity expressed by XTT of *C. albicans* grown for 48 hours in the absence or presence of CAS 0 to 1 µg/ml or BRI 20 µM+CAS 0 to 1 µg/ml. Percentage of survival expressed as colony forming units/ml of *C. albicans* cells grown for 48 hs in the absence or presence of CAS 0 to 1 µg/ml or BRI 20 µM+CAS 0 to 1 µg/ml. (C) Metabolic activity expressed by XTT of *C. albicans* CAS-resistant strains grown for 48 hours in the absence or presence of CAS 0.5 µg/ml, BRI 5 to 20 µM or BRI 5 to 20 µM+CAS 0.5 µg/ml. (D) Metabolic activity expressed by XTT of *C. auris* grown for 48 hours in the absence or presence of CAS 0.125 µg/ml, BRI 10 µM or BRI 10 µM+CAS 0.125 µg/ml. Metabolic activity expressed by XTT of *C. auris* 467/2015 strain grown for 48 hours in the absence or presence of CAS 0 to 1 µg/ml or BRI 10 µM+CAS 0 to 1 µg/ml. Percentage of survival expressed as colony forming units/ml of *C. auris* 467/2015 strain grown for 48 hs in the absence or presence of CAS 0 or 1 µg/ml or BRI 10 µM+CAS 0 to 1 µg/ml. (*, *p* < 0.05, **, *p <* 0.01, ***, *p* < 0.001, and ****, *p* < 0.0001). All the results are the average of three repetitions ± standard deviation.

**Table 2.**
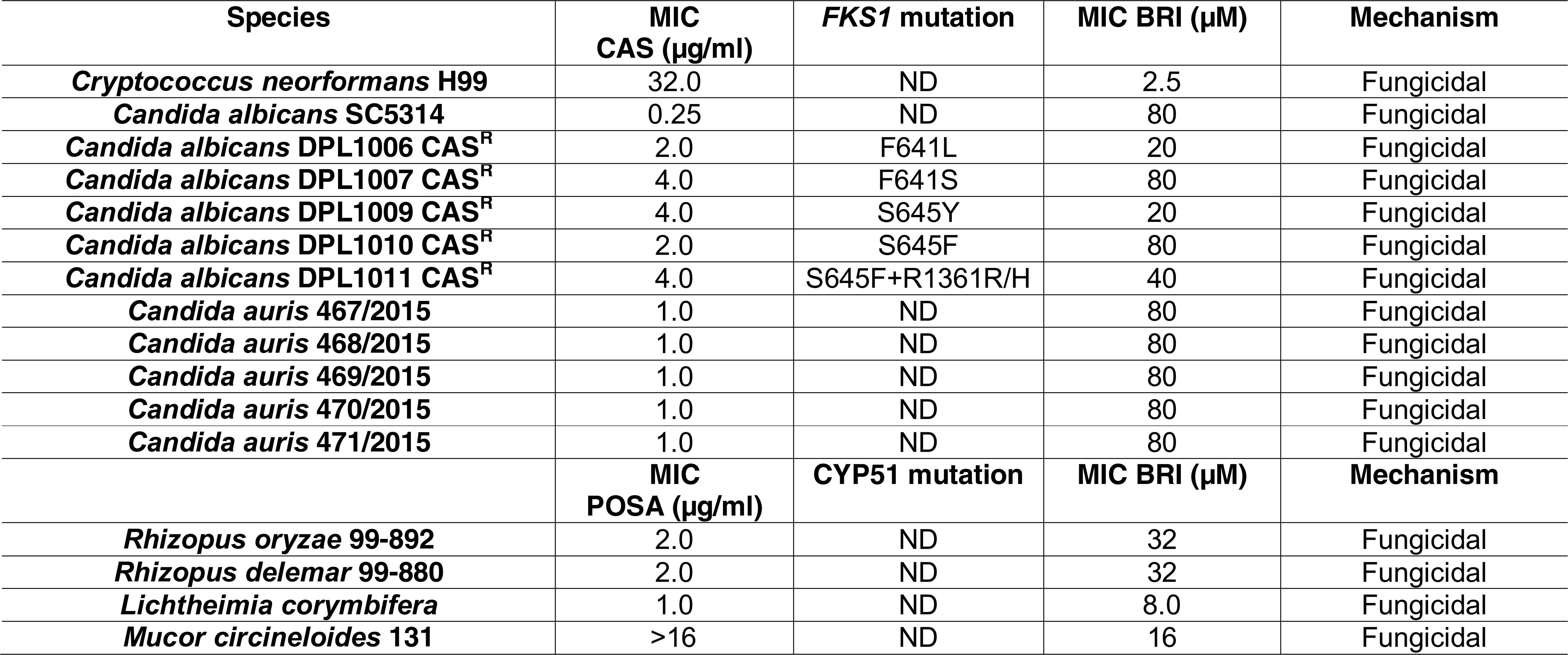
Brilacidin MICs for human pathogenic fungi.

Taken together, these results indicate that BRI is able to potentiate CAS activity for different human fungal pathogens, including *C. neoformans.* Interestingly, *C. neoformans* is very sensitive to BRI alone and BRI is fungicidal against this fungus. Thus, BRI is a novel therapeutic against *C. neoformans* alone or in combination with CAS since it potentiates the latter’s activity into a fungicidal drug. BRI is also able to convert CAS into a fungicidal drug in *C. auris*.

### BRI enhances the activity of POSA against Mucorales fungi

POSA is used for treating patients with lethal mucormycosis as a step-down therapy to liposomal amphotericin B (LAMB) or in lieu of LAMB in patients who are refractory or intolerate to the latter drug (Cornely *et al*. 2019). We investigated if BRI could potentiate the activity of POSA against the most common causes of mucormycosis including *Rhizopus delemar, R. oryzae, Lichtheimia corymbifera, Mucor circineloides.* The growth of these fungi was measured in a checkerboard 96-well assay for which BRI concentrations of 64-0.12 µg/mL and POSA concentrations of 16-0.25 µg/mL were used. In two independent experiments, the Fractional Inhibitory Concentrations (FIC) indices for these two drugs ranged between 0.51-1.12 indicating additive effect for BRI when added to POSA against the tested Mucorales fungi (Table 3). Thus, unlike VOR and BRI against *A. fumigatus,* it appears that BRI can potentiate the activity of POSA against Mucorales fungi *in vitro*.

**Table 3.**
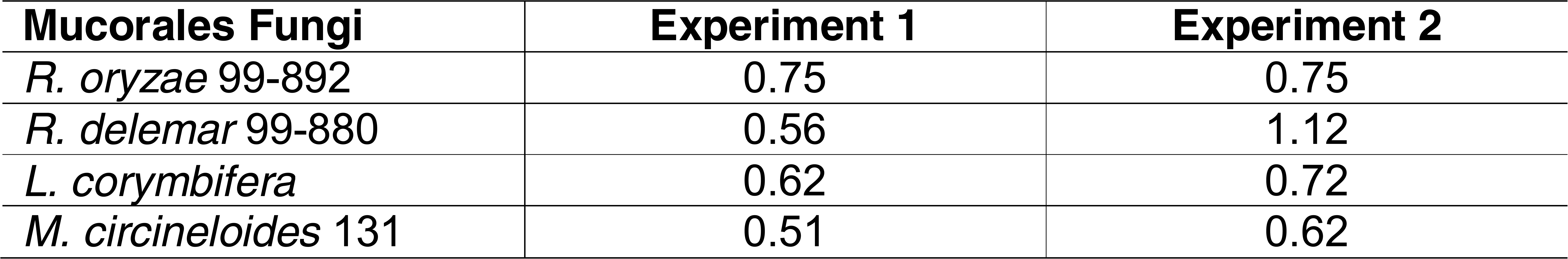
FIC indices of BRI + POSA combination against Mucorales fungi.

### BRI combined with CAS is not toxic to human cells and decreases the *A. fumigatus* fungal burden in a chemotherapeutic murine model

Toxicity assessment of brilacidin in A549 pulmonary cells was initially performed by incubating the cells either with 40 or 80 µM of BRI with or without increasing CAS concentrations for 48 h, after which cell viability was assessed by XTT assay (Figure 5A). Neither BRI, CAS, or their combinations reduced cell viability when compared to the DMSO vehicle control (Figure 5A). We also compared the ability of the combination of BRI+CAS to the standard of care, VOR, in controling *A. fumigatus* cell growth when infecting A549 pulmonary cells. While VOR, CAS, and BRI monotherapy killed ∼ 60%, 25%, and 0%-10%, respectively, a combination of BRI (20-80 µM) + CAS (100 µg/ml) resulted in 50-85% *A. fumigatus* killing (Figure 5B).

**Figure 5.**
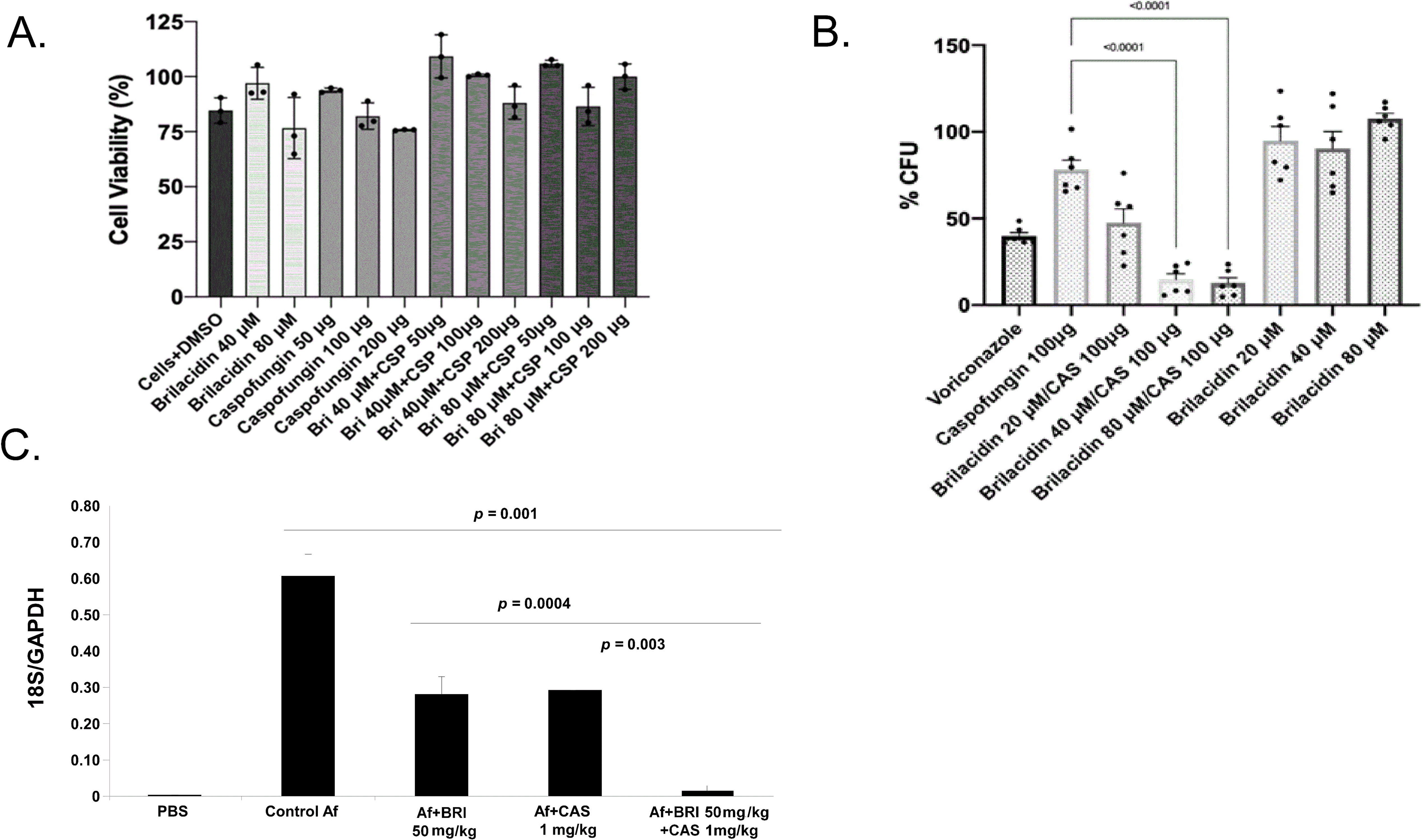
The combination of BRI+CAS is not toxic to human cells and can significantly decrease the *A. fumigatus* fungal burden in a chemotherapeutic murine model. (A) A549 lung cells grown in the absence or presence of different concentrations of BRI and CAS. Percentage of cell viability is expressed as the absorbance value of experiment well/absorbance value of control well x 100. All the results are the average of three repetitions ± standard deviation. (B) A549 lung cells were infected with *A. fumigatus* conidia in the presence or absence of different concentrations of BRI and CAS. Percentage of cell viability is expressed as the absorbance value of experiment well/absorbance value of control well x 100. All the results are the average of three repetitions ± standard deviation. (C) Fungal burden was determined 72 hours post-infection by real-time qPCR based on 18S rRNA gene of *A. fumigatus* and an intronic region of the mouse GAPDH gene. Fungal and mouse DNA quantities were obtained from the Ct values from an appropriate standard curve. Fungal burden was determined through the ratio between ng of fungal DNA and mg of mouse DNA. The results are the means (± standard deviation) of five lungs for each treatment. Statistical analysis was performed by using *t*-test.

Finally, we investigated if BRI+CAS could impact *A*. *fumigatus* virulence in a chemotherapeutic murine model of IPA. Fungal burden in the lungs was approximately 50% reduced after 3 days post-infection in mice treated either with CAS (1mg/kg) or BRI (50 mg/kg) when compared with the non-treated mice (Figure 5C). However, the combination of BRI+CAS significantly reduced the fungal burden by ∼95% when compared with the non-treated mice (Figure 5C). These results strongly indicate that the combination BRI+CAS is able to clear *A*. *fumigatus* infection in the lungs in a chemotherapeutic murine model of IPA.

Taken together, these data indicate that the combination treatment of BRI+CAS is non-toxic to mammalian cells *in vitro* and is able to enhance clearance of *A. fumigatus* infection in pulmonary cells *in vitro* and *in vivo* when compared to monotherapy alone.

## Discussion

Due to the reduced number of available antifungal agents and the increased emergence of antifungal resistance in the clinical environment, there is an urgent need for novel antifungal drugs. There are few new potential antifungal drugs at different stages of clinical development, such as fosmanogepix (a novel Gwt1 enzyme inhibitor), ibrexafungerp (a novel oral glucan synthase inhibitor), olorofim (a novel dihyroorotate dehydrogenase enzyme inhibitor), opelconazole (a novel triazole optimized for inhalation), and rezafungin (an echinocandin with an extended half-life) (Hoenigl *et al*., 2021). An alternative approach to the development of novel antifungal drugs is the repositioning or repurposing of existing drugs. This approach is particularly promising because usually it identifies drugs that can synergize or enhance the activity of current antifungal agents and in many cases overcome developed resistance to a certain class of antifungals. Here, we have screened four repurposing libraries and identified enhancers and synergizers of CAS activity. We concentrated our attention on one of these synergizers, BRI, that is able to synergize not only with CAS but also VOR.

Human host defense peptides (HDPs or antimicrobial peptides), one class of which are the human defensins, are part of the innate immune response (Kuroda and Caputo, 2013; Sgolastra *et al*., 2013). BRI is a polymer-based, non-peptidic small molecule drug candidate exhibiting antimicrobial and immunomodulatory properties, designed to mimic the structure and activity of HDPs (Scott and Tew, 2017; Matos de Opitz and Sass, 2020). BRI has been tested in several clinical trials, such as (i) treatment for acute bacterial skin and skin structure infections (ABSSSI) caused by *Staphylococcus aureus* (Clinical Trials NCT01211470, https://www.clinicaltrials.gov/ct2/show/NCT01211470?term=polymedix; (Bassetti et al., 2020; Mensa et al., 2014), (ii) oral mucositis (NCT02324335, https://clinicaltrials.gov/ct2/show/NCT04784897), and more recently (iii) Covid-19/SARS-Cov-2 (https://clinicaltrials.gov/ct2/show/NCT04784897; (Bakovic et al., 2021; Hu et al., 2022). Like other antimicrobial peptides, BRI can depolarize the *A. fumigatus* cell membrane (Sahl *et al*., 2005). However, it is not clear from our studies if BRI is localized on the cell surface, or if it can be transported intracellularly and interacts with other cellular proteins. We investigated how BRI can convert CAS into a fungicidal drug by a combination of screening PKIs and a collection of *A. fumigatus* null mutants for the catalytic subunits of phosphatases. We observed two PKI compounds, a p21-Activated Kinase Inhibitor FRAX486 and a STK25 inhibitor PP121, both members of the sterile 20 superfamily of kinases, as potentiating the BRI activity against *A. fumigatus*. Since p21 activated kinases (PAKs) are involved in signaling through MAPK pathways (Boyce and Andrianopoulos, 2011), we tested if other *A. fumigatus* MAPK are more sensitive to BRI and identified MpkA as being so, the apical MAPK that is regulated by protein kinase C and coordinates the CWI pathway (Valiante *et al*., 2015). Complementary to this screening, we also identified calcineurin as more sensitive to BRI and increased CrzA:GFP translocation to the nucleus in the presence of BRI+CAS. Since Δ*mpkA* and Δ*calA* have reduced growth when compared to the wild-type strains, these results were confirmed by using chelerenthrine and calphostin C (inhibitors of protein kinase C) and cyclosporin (an inhibitor of calcineurin). These results suggest that BRI is inhibiting the *A. fumigatus* CWI pathway suppressing the compensatory reactions, such as replacement of β-1,3-glucan by chitin or further remodelling of the cell wall, hallmarks of the CAS fungistatic effect. Thus, we propose that BRI is able to convert CAS into a fungicidal drug through multiple mechanisms of action, encompassing functional changes/depolarization of the microorganism cell membrane, interference to calcineurin signaling, and inhibition of the cell wall integrity pathway and necessary compensatory reactions (since CAS is also present and prevents further β-1,3-glucan biosynthesis). Consequently, cell wall remodelling, which prevents resistance against osmotic forces, is inhibited which then leads to cell lysis. Alterations in the β-1,3-glucan synthase lipid environment could also be important (Satish *et al*., 2019). Recently, a novel off-target mechanism of action for CAS was proposed in which CAS-induced ROS alters the lipid composition around the β-1,3-glucan synthase, changing its conformation and making it less sensitive to CAS (Satish *et al*., 2019). It is possible that BRI also influences the lipid composition of the cell membrane (*e.g.,* by interacting with ergosterol) making the β-1,3-glucan synthase more accessable to CAS. It is likely that BRI can potentiate VOR activity through the disorganization of the cell membrane which would not allow the proper deposition of ergosterol leading to impaired function of the sterol. An important feature of BRI was its ability to overcome CAS-resistance in both *A. fumigatus* and *C. albicans* but not VOR-resistance in *A. fumigatus*. The molecular basis for CAS-resistance in both species is related to hotspot mutations in the *fks1* gene that encodes β-1,3-glucan synthase (Perlin, 2011). All the above suggested mechanisms could also contribute to the bypass of CAS-resistance by BRI. It remains to be investigated how BRI can potentiate VOR, and how BRI can bypass CAS-resistance in *A. fumigatus*. Equally important is the finding that BRI can enhance the activity of POSA against Mucorales fungi which cause mucormycosis, a disease with high mortality rates of 50-90%, thereby suggesting a potential enhancement in the therapeutic outcome of this lethal disease.

A remarkable observation of our work is the fact that BRI can potentiate CAS activity not only against *A. fumigatus*, but also *C. albicans*, *C. auris*, and *C. neoformans*. There are very few therapeutic options against the treatment of cryptococcosis (Iyer *et al*., 2021) and BRI demonstrated activity against *C. neoformans* in low concentrations (MIC=2.5 µM). *C. neoformans* is intrinsically resistant to CAS and only very high non-physiological CAS concentrations can partially inhibit *C. neoformans* growth (Johnson and Perfect, 2003). However, *C. neoformans* β-1,3-glucan synthase is very sensitive to CAS (Maligie and Selitrennikoff, 2005), which suggests that other mechanisms unrelated to β-1,3-glucan synthase resistance are important for CAS resistance. We have not investigated the molecular basis for BRI potentiation of CAS in *C. neoformans*. Nevertheless, there is a report showing that cyclosporine and FK506 (tacrolimus) can enhance the activity of caspofungin against *C. neoformans* (Del Poeta *et al*., 2000), suggesting the possibility that the same mechanism observed for *A. fumigatus* is also acting in *C. neoformans.* Interestingly, amphotericin B which disturbs the cell membrane via self-assembly into an extramembranous sponge that rapidly extracts ergosterol from fungal membranes, synergizes with CAS against *C. neoformans* (Banerjee *et al*., 2016; Guo *et al*., 2021) and Mucorales fungi (Spellberg *et al*. 2005). This suggests that an analogous mechanism of cell membrane disruption by amphotericin could enhance CAS actitivity in *C. neoformans*, reinforcing the notion that BRI is impacting the organization of the cell membrane in this species.

Initial studies demonstrated that the combination of BRI+CAS can significantly reduce the *A. fumigatus* fungal burden in an immunosuppressed murine model, strongly indicating the therapeutic potential of BRI + CAS against IPA. Further studies will address possible formulations to deliver BRI and CAS together and to understand the pharmacokinetics. Our results demonstrate the promise of repurposing studies and the screening of well annotated compounds for potentiating the activity of current antifungal drugs.

## Materials and Methods

### Strains, media and cultivation methods

The *A. fumigatus*. *Candida spp., C. neoformans* strains and Mucorales fungi used in this work are listed in Supplementary Table S4. *Aspergillus* strains were grown in minimal medium (MM; 1% [wt/vol] glucose, 50 mL of 20x salt solution, trace elements, 2% [wt/vol], pH 6.5. For solid minimal medium 2% agar was added) at 37°C. Solutions of trace elements and salt solution are described by Käfer et al., 1977. For the animal studies, *A. fumigatus* strain (wild-type, CEA17) was grown on MM. Fresh conidia were harvested in PBS and filtered through a Miracloth (Calbiochem). Conidial suspensions were spun for 5 min at 3,000x g, washed with PBS, counted using a hemocytometer and resuspended at a concentration of 5.0 x 10^7^ conidia/ml. *Candida spp.* and *C. neoformans* strains were grown and maintained on YPD (1% yeast extract, 2% peptone and 2% glucose). For Mucorales fungi, all organisms were grown on potato dextrose agar (PDA; BD Diagnostic, New Jersey) plates by sprinkling silica beads that have been absorbed with spore suspension of each organism. The plates were incubated at 37°C for 4-7 days. Spores were collected in endotoxin-free Dulbecco phosphate buffered saline (PBS) containing 0.01% or 0.2% Tween 80 for Mucorales or *Aspergillus*, respectively. Collected spores were washed with PBS, and counted with a hemocytometer to prepare the final inocula.

### Library drug screenings

The Pandemic Response box (400 compounds) and COVID box (160 compounds) (both available at https://www.mmv.org), the National Institutes of Health (NIH) clinical collection (727 compounds) (available at https://pubchem.ncbi.nlm.nih.gov/source/NIH%20Clinical%20Collection) and the SGC’s epigenetic chemical probe and SGC’s pharma donated chemical probe libraries (115 compounds, see https://www.thesgc.org/chemical-probes) totaling 1,402 compounds, were screened in the current study. The primary screening was performed against the *A. fumigatus* wild-type strain by using the chemical libraries diluted in dimethyl sulfoxide (DMSO). The ability of each compound in combination (or not) with caspofungin (CAS) in blocking the fungal growth was visually determined. Briefly, each well of a flat-bottom polystyrene microplate was filled with 198 µL of liquid MM containing 1 x 10^4^ conidia/mL of *A. fumigatus* (wild-type strain). Subsequently 20 µM of each chemical compound was added in combination (or not) with 0.2 µg/mL of CAS to each well. This concentration represents the MEC for CAS against *A. fumigatus*. Plates were statically incubated for 48 h at 37°C. Wells containing only medium, CAS [0.2 µg/mL] or DMSO were used as controls. Compounds presenting over 80% of visual fungal growth inhibition (in combination or not with CAS) were selected for further studies. All experiments were done in triplicate.

### Alamar blue assays

The inhibition of the metabolic activity of *A. fumigatus* triggered by the drugs selected in the first screening was assessed by using Alamar blue (Life Technologies) according to Yamaguchi et al. (2002). The experiment was done by inoculation of 100 µL of liquid MM containing 2.5 x 10^3^ conidia/mL of the *A. fumigatus* wild-type strain supplemented or not with CAS [0.2 µg/mL] plus increasing concentration of each selected drug [0.6 to 20 µM] and 10% Alamar blue in 96-well plates. As positive controls, the drugs were replaced by the same volume of the medium. As the negative control, wells were filled with 90 µL of liquid MM plus 10 µL of Alamar Blue. Plates were incubated for 48 h at 37°C without shaking and results were read spectrophotometically by fluorescence (570 nm excitation– 590 nm emission) in a microplate reader (SpectraMax® Paradigm® Multi-Mode Microplate Reader; Molecular Devices). Enhancers were defined as compounds that alone inhibited over 30% of *A. fumigatus* metabolic acitivity but in combination with CAS inhibited even more, while synergizers were defined as compounds which alone inhibited less than 30 % of the fungal metabolic activity but in combination with CAS inhibited more than 30 %.

A protein kinase inhibitors (PKI) library was also screened in combination with BRI. In total, 58 PKI were analyzed. Briefly, 100 µL of liquid MM containing 2.5 x 10^3^ conidia/mL of the *A. fumigatus* wild-type strain plus 10% alamar blue was inoculated with increasing concentration of PKI [5-80 µM] in the presence (or not) of BRI [20 µM] and incubated 48 h at 37°C without shaking. To analyze the sensitivity of the null mutants Δ*calA*, Δ*mpkA* and their complementing strains to BRI, 100 µL of liquid MM containing 2.5 x 10^3^ conidia/mL of each strain was inoculated in the presence (or not) of BRI [20 µM] plus 10% alamar blue and incubated 48 h at 37°C without shaking. To check if cyclosporine (CsA), chelerenthrine and/or calphostin C synergize with BRI, variable concentrations of each one of these drugs was analyzed in the presence of BRI [20 µM]. A total 100 µL of liquid MM containing 2.5 x 10^3^ conidia/mL of the *A. fumigatus* wild-type strain plus 10% alamar blue was inoculated with increasing concentrations of CsA [25-200 µg/mL], chelerenthrine [0.78-6.25 µg/mL] and calphostin C [6.25-50 µg/mL] in the presence (or not) of BRI [20µM]. Plates were incubated for 48 h at 37°C without shaking. All experiments containing alamar blue were, after 48h incubation, read spectrophotometically by fluorescence and analyzed as previously described. Experiments were repeated at least three times.

### Minimal inhibitory concentration (MIC)

The BRI drug used for MIC assays was kindly supplied by Dr. William DeGrado and solubilized in DMSO. The minimal inhibitory concentration (MIC) of BRI for *A. fumigatus* or Mucorales fungi was determined based on the M38-A2 protocol of the Clinical and Laboratory Standards Institute (CLSI 2008) and for yeasts using M27-A3 method (CLSI, 2017). In brief, the MIC assay was performed in 96-well flat-bottom polystyrene microplate where 200 µL of a suspension (1 x 10^4^ conidia/mL) prepared in liquid MM was dispensed in each well and supplemented with increasing concentration of BRI (ranging from 0 to 160 µM). Plates were incubated at 37°C without shaking for 48h and the inhibition of growth was evaluated. The MIC was defined as the lowest drug concentration that visually attained 100% of fungal growth inhibition compared with the control well. Wells containing only MM and DMSO were used as a control. Similar protocol was used for yeast organisms, except that we used RPMI-1640, 1 x 10^3^ cells/mL/well and incubated the plates for 48h (*Candida* spp.) or 72h (*C. neoformans*).

### FIC index analysis

To determine synergy, indifferent, or antagonism between and geldanamycin and BRI (against *A. fumigatus*) and BRI and POSA (against Mucorales fungi), we used the FIC index method (Meletiadis et al. 2005). Briefly, for all of the wells of the microtitration plates that corresponded to an MIC, the sum of the FICs (ΣFIC) was calculated for each well with the equation ΣFIC = FICA + FICB = (CA(comb)/MICA(alone)) + (CB(comb)/MICB(alone)), where MICA(alone) and MICB(alone) are the MICs of drugs A and B when acting alone, and CA(comb) and CB(comb) are the concentrations of the drugs A and B at the iso-effective combinations. An FIC index of < 0.5 indicates synergism, > 0.5–1 indicates additive effects, > 1 to < 2 indifference, and ≥ 2 is considered to be antagonism (Faleiro et al. 2013).

### Combination of BRI and CAS against yeasts

For measuring the effect of the combination BRI+CAS against yeast fungal pathogens, two methods were used: (i) metabolic activity by XTT-assay as described by Bastos *et al*. (2019) and (ii) colony forming units (CFUs). For the first method, *C. neoformans*, *C. albicans*, and *C. auris* 10^4^ cells were inoculated in RPMI-1640 supplemented with CAS 0 to 32 µg/ml (for *C. neoformans*) and 0 to 1 µg/ml (for *Candida spp.*) or the same concentrations of CAS combined with BRI 0.625 µM (for *C. neoformans*), BRI 20 µM (for *C. albicans*), and BRI 10 µM (for *C. auris*). After 48 h of incubation, the viable cells were revealed using XTT-assay as described by Bastos et al. (2019).

XTT-assays were also used for *C. albicans* caspofungin resistant strains but with CAS 0.5 µg/ml combined with BRI 5, 10 or 20 µM. The same experimental design was used for the CFUs determination, except that after 48 hs the cells present in the wells were plated on YPD (yeast extract 10g, peptone 20g, dextrose 20g, agar 20g, water 1000 ml) and the plates were incubated at 30°C for 24-48h for determining the survival percentage. The results are the average of three repetitions and are expressed as average ± standard deviation.

### Conidial viability exposed to brilacidin (BRI), voriconazole (VOR) and caspofungin (CAS)

The viability of *A. fumigatus* conidia exposed to CAS+BRI or VOR+BRI was assessed by plating the cells after being treated. Initially, a suspension containing 1 x 10^4^ conidia/mL of *A. fumigatus* cells was prepared in liquid MM and 200 µL of this suspension was dispensed in each well of a 96-well polystyrene microplate supplemented with CAS [0.2 or 0.50 µg/mL] + BRI [20µM] or VOR [0.125 or 0.25 µg/mL] + BRI [20 µM]. After 48h incubation at 37°C, a total of 100 conidia was plated in solid complete medium (YAG) [2% (w/v) glucose, 0.5% (w/v) yeast extract, trace elements] or minimal medium [1% (w/v) glucose, nitrate salts, trace elements, pH 6.5] and let to grow at 37°C for 36h. The number of viable colonies was determined by counting the number of colony-forming units (CFUs) and expressed in comparison with the negative control (no germinated and untreated conidia), which gives 100% survival. Results are expressed as means and standard deviations (SD) from three independent experiments.

### Biofilm assay

To test the susceptibility of pre-formed *A. fumigatus* biofilms to VOR, CAS and to the combination of CAS + BRI and VOR + BRI, a suspension containing 10^6^ conidia per mL of the wild-type strain (CEA17) was prepared in liquid MM and 100 µL of it was inoculated in each well of a 96-well plate. After 24 h of incubation at 37°C, 50 µL of fresh MM containing CAS, VOR or the combination of VOR and CAS with BRI was added to the biofilm to reach the final concentration as indicated and incubated for a further 12 h at 37°C. Wells containing untreated conidia were used as a positive control. After, the metabolic activity of the cells was evaluated by adding 50 µL of an aqueous XTT solution (1mg/mL of XTT and 125 µM of menadione) to each well. The plate was incubated for an additional 1h at 37°C, centrifuged (2000 rpm, 5 minutes) and 100µL of the supernatant was transferred to a flat-bottomed 96-well plate. The absorbance was measured at 450 nm on a plate reader (Synergy HTX Multi-Mode Reader-BioTek Instruments). The XTT assays were performed in six replicates.

### Phosphatase and kinase null mutant screening

An *A. fumigatus* phosphatase deletion library encompassing 25 null mutants for phosphatase catalytic subunits (Winkelströter et al., 2015) was screened for sensitivity to the combination of CAS + BRI. *A. fumigatus* null mutants for MAPK (Δ*sakA*, Δ*mpkC*, Δ*sakA*; Δ*mpkC*, Δ*mpkB* and Δ*mpkA)* were also screened. The assay was performed in a 96-well flat-bottom polystyrene microplate. In each well a total of 200 µL of liquid MM plus conidia from the different mutants (1 x 10^4^ conidia/mL) was incubated in the presence of BRI [20 µM]. Plates were incubated at 37°C without shaking for 48h and the inhibition of growth was visually evaluated. Wells containing only MM and DMSO were used as a control.

### Membrane potential determination

The effect of the CAS [0.125 µg/mL], BRI [1 µM] or the combination CAS + BRI ([0.125 µg/mL] and [1 µM], respectively) on the cell membrane potential was assessed by using the bis-(1,3-dibutylbarbituric acid) trimethine oxonol - DiBAC4(3) reagent (Invitrogen, Carlsbad, CA, USA) according to Veerana et al., 2021 with modifications. *A. fumigatus* conidia were inoculated on coverslips in 5mL of liquid MM and cultivated for 16h at 30°C. Further, coverslips containing adherent germlings were left untreated or treated with CAS, BRI or CAS + BRI plus 3 µg/ml DIBAC_4_(3) and incubated for 30 minutes at 30°C in the dark. After, the germlings were washed with sterile PBS (140mM NaCl, 2 mM KCl, 10mM NaHPO4, 1.8 mM KH2 PO4, pH 7.4). The fluorescence was analyzed with excitation wavelength of 450nm-490nm, and emission wavelength of 525 nm – 550 nm on the Observer Z1 fluorescence microscope (Carl Zeiss) using the 100x with Differential interference contrast (DIC) images. Fluorescent images were captured with an AxioCam camera (Carl Zeiss, Inc.) and processed using the AxioVision software (version 4.8). In each experiment, at least 50 germlings were counted and the experiment repeated at least 3 three times.

### Cytotoxicity assay

Cytotoxicity assays in A549 human lung cancer cells were performed using XTT assay as indicated in the manufacturers’ instructions. Cells (2X10^5^ cells/well) were seeded in 96-well tissue plates and incubated in Dulbecco’s Modified Eagle Medium (DMEM) culture medium. After 24 h of incubation, the cells were treated with BRI (40 and 80 µM/well), CAS (50, 100 and 200 µg/well) or in different CAS+BRI combinations. After 48 h incubation, the cell viability was assessed by using the XTT kit (Roche Applied Science) according to the manufacturer’s instructions. Formazan formation was quantified spectrophotometrically at 450 nm (reference wavelength 620 nm) using a microplate reader. The experiment was made in three replicates. Viability was calculated using the background-corrected absorbance as follows: Cell viability (%) = absorbance value of experiment well/absorbance value of control well x 100.

### Killing assay

The type II pneumocyte cell line A549 was cultured using DMEM (ThermoFischer Scientific, Paisley, UK) supplemented with 10% fetal bovine serum (FBS) and 1% penicillin–streptomycin (Sigma-Aldrich, Gillingham, UK) and seeded at a density of 10^6^ cells/ml in 24-well plates (Corning). The cells were treated with Brilacidin (20, 40 and 80 µM/well), Caspofungin (100 µg/well) or in different combinations between them and challenged with *A. fumigatus* conidia at a multiplicity of infection of 1:10. After 24 h of incubation in 5% CO_2_ the culture media was removed, and 2 ml of sterile water was added to the wells. A P1000 tip was then used to scrape away the cell monolayer and the cell suspension was collected. This suspension was then diluted 1:1000 and 100 μl was plated on Sabouraud Dextrose Agar Media before the plates were incubated a 37°C overnight. The numbers of CFUs were determined after 24 h of growth. A volume of 50 μl of the inoculum adjusted to 10^3^/ml was also plated on SAB agar to correct CFU counts. The CFU percentage for each sample was calculated and the results were plotted using Graphpad Prism (GraphPad Software, Inc., La Jolla, CA, USA). A *p* value ≤0.001 was considered significant.

### Fungal burden

Inbred female mice (BALB/c strain; body weight, 20–22 g) were housed in vented cages containing five animals. Mice were immunosuppressed with cyclophosphamide (150 mg/kg of body weight), which was administered intraperitoneally on days -4, -1 and 2 prior to and post infection (infection day is “day 0”). Hydrocortisonacetate (200 mg/kg body weight) was injected subcutaneously on day -3. Mice (5 mice per group) were anesthetized by halothane inhalation and infected by intranasal instillation of 20 µL of 1.0 x 10^6^ conidia of *A. fumigatus* CEA17(wild-type) (the viability of the administered inoculum was determined by incubating a serial dilution of the conidia on MM medium, at 37°C). As a negative control, a group of 5 mice received PBS only. On the same day of infection (day 0), mice received concomitantly the first dose of treatment with BRI (50mg per kg of body weight) and/or CAS (1mg per kg of body weight), administered intraperitoneally. The second dose of drugs was administered 24 hours after infection. Animals were sacrificed 72 h post-infection, and the lungs were harvested and immediately frozen in liquid nitrogen. Samples were lyophilized and homogenized by vortexing with glass beads for 5 min, and DNA was extracted via the phenol/chloroform method.

DNA quantity and quality were assessed using a NanoDrop 2000 spectrophotometer (Thermo Scientific). Quantitative real-time PCRs were performed using 400 ng of total DNA from each sample, and primers to amplify the 18S rRNA region of *A. fumigatus* and an intronic region of mouse GAPDH (glyceraldehyde-3-phosphate dehydrogenase). Six-point standard curves were calculated using serial dilutions of gDNA from *A. fumigatus* strain and the uninfected mouse lung. Fungal and mouse DNA quantities were obtained from the threshold cycle (Ct) values from an appropriate standard curve.

### Statistical analysis

Grouped column plots with standard deviation error bars were used for representations of data. For comparisons with data from wild-type or control conditions, we performed one-tailed, paired *t* tests or one-way analysis of variance (ANOVA). All statistical analyses and graphics building were performed by using GraphPad Prism 5.00 (GraphPad Software).

## Acknowledgements

We thank the Fundação de Amparo à Pesquisa do Estado de São Paulo (FAPESP) grants numbers 2016/07870-9 and 2021/04977-5 (G.H.G.) and the Conselho Nacional de Desenvolvimento Científico e Tecnológico (CNPq) grant numbers 301058/2019-9 and 404735/2018-5 (G.H.G.), both from Brazil, and the National Institutes of Health/National Institute of Allergy and Infectious Diseases grants R01AI153356, and R01AI063503 to ASI from the USA. A special thanks to CAPES for providing the grant for the postdoctoral position for Pedro F. N. Souza. The epigenetic and donated chemical probe libraries were supplied by the Structural Genomics Consortium under an Open Science Trust Agreement: http://www.thesgc.org/click-trust.

