## Supplementary figures and images for "A host defense peptide mimetic, brilacidin, potentiates caspofungin antifungal activity against human pathogenic fungi"

### Supplementary Figure S1

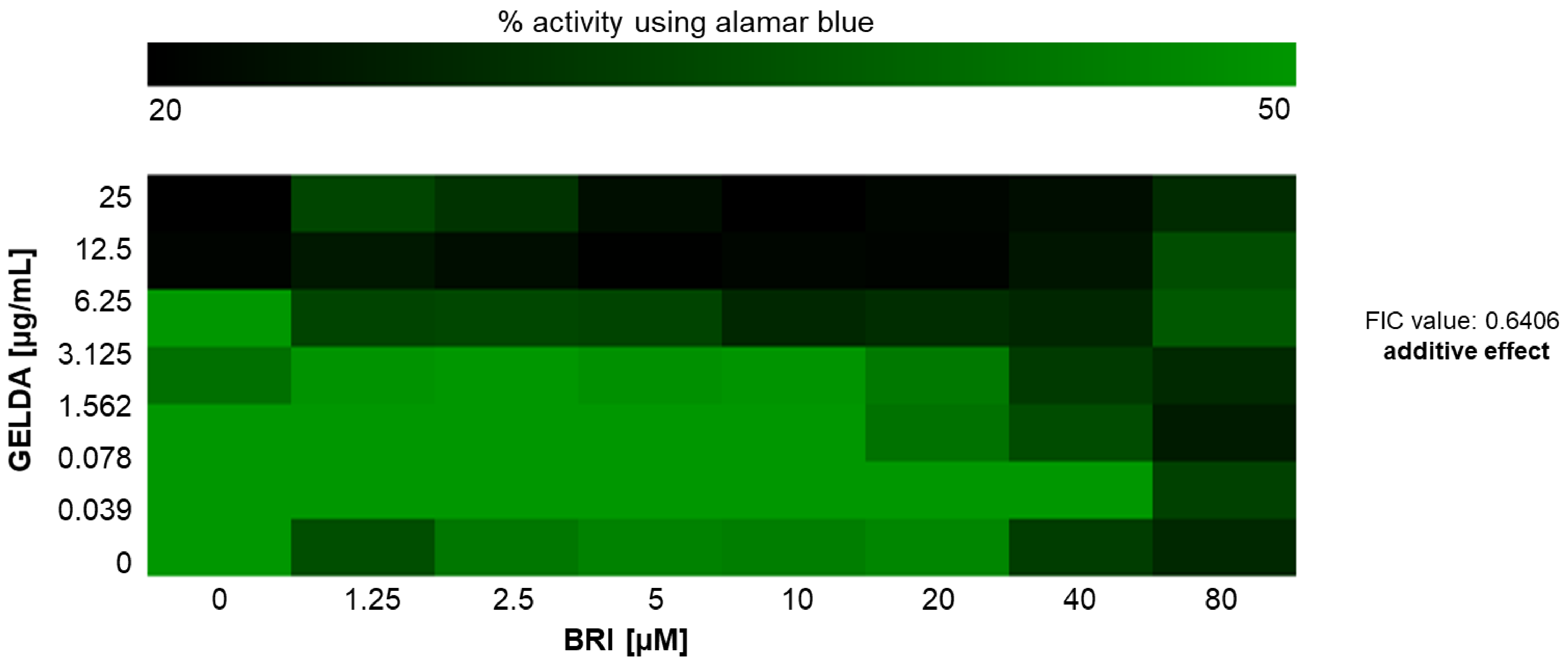


Supplementary Figure S1 - Checkerboard for geldanamycin and brilacidin against *A. fumigatus*
