## Supplementary Table S1 for "A host defense peptide mimetic, brilacidin, potentiates caspofungin antifungal activity against human pathogenic fungi"

Supplementary Table S1 - BRI+VOR cannot overcome *A. fumigatus* VOR-resistance

|  |  |  |  | **VOR [µg/ml]** | | **BRI [µM]** | **0.5 µg/ml VOR** | | **2 µg/ml VOR** | |
| --- | --- | --- | --- | --- | --- | --- | --- | --- | --- | --- |
| **Strain** | **Mechanism of**  **VOR^R^** | **MIC VOR [µg/ml]** | **MIC BRI [µM]** | **0.5** | **2** | **20** | **20 µM BRI** | **40 µM BRI** | **20 µM BRI** | **40 µM BRI** |
| **WT (CEA17)** | WT | 2 | 80 | --- | --- | + | --- | --- | --- | --- |
| **CYP-15-184** | TR34/L98H | 8 | 80 | + | + | + | + | + | + | + |
| **CYP-15-190** | TR34/L98H | 8 | 80 | + | + | + | + | + | + | + |
| **CYP-15-192** | TR34/L98H | 4 | 80 | + | + | + | + | + | + | + |
| **CYP-15-195** | TR34/L98H | 8 | 80 | + | + | + | + | + | + | + |
| **CYP-15-202** | TR34/L98H | 8 | 80 | + | + | + | + | + | + | + |
| **CYP-15-212** | TR34/L98H | 8 | 80 | + | + | + | + | + | + | + |
| **CYP-15-213** | TR34/L98H | 4 | 80 | + | + | - | + | + | + | + |
| **CYP-15-215** | Unknown | 8 | 80 | + | + | - | + | + | + | + |
| **CYP-15-220** | TR34/L98H | 4 | 80 | + | + | + | + | + | + | + |
| **CYP-15-221** | TR34/L98H | 8 | 80 | + | + | + | + | + | + | + |
| **CYP-15-222** | TR34/L98H | 4 | 80 | + | + | + | + | + | + | + |
| **CYP-15-224** | Unknown | 8 | 80 | + | + | + | + | + | + | + |
| **CYP-15-225** | Unknown | 8 | 80 | + | + | + | + | + | + | + |
| **CYP-15-226** | TR34/L98H | 4 | 80 | + | + | + | + | + | + | + |
| **CYP-15-228** | TR34/L98H | 4 | 80 | + | + | + | + | + | + | + |
| **CYP-15-229** | TR34/L98H | 8 | 80 | + | + | + | + | + | + | + |
| **CYP-15-230** | TR34/L98H | 8 | 80 | + | + | + | + | + | + | + |
| **CYP-15-231** | TR34/L98H | 4 | 80 | + | + | + | + | + | + | + |
| **CYP-15-75** | Unknown | 8 | 80 | + | + | + | + | + | + | + |
| **CYP-15-108** | Unknown | 8 | 80 | + | + | + | + | + | + | + |
| **CYP-15-109** | Unknown | >8 | 80 | --- | --- | + | --- | --- | --- | --- |
| **CYP-15-147** | Unknown | 8 | 80 | + | + | + | - | - | + | + |
| **F14946** | Unknown | 8 | 80 | + | + | + | + | + | + | + |
| **20089320** | Unknown | 8 | 80 | + | + | + | + | + | + | + |

Obs.: 1) VOR=voriconazole; BRI=Brilacidin. (+): no inhibition; (-): partial inhibition; and (---) total inhibition;

2) In the experiment, one strain (CYP15-15-109) was totally inhibited by VOR, showing Wild Type response; also, one strain (CYP15-15-147) was partially inhibited by VOR 0.5 µg/ml+BRI but the inhibition was not seen at VOR 2.0 µg/ml+BRI.
